# Complementary BOLD- and ADC-fMRI explore the role of lateral superior colliculus in flicker fusion frequency

**DOI:** 10.1101/2025.07.24.666486

**Authors:** Jean-Baptiste Pérot, Andreea Hertanu, Arthur Spencer, Jasmine Nguyen-Duc, Nikolaos Molochidis, Valerio Zerbi, Maxime Yon, Ileana Jelescu

## Abstract

The transition from static to dynamic vision is encoded in the superior colliculus, as recently shown using blood-oxygen-level-dependent functional MRI (BOLD-fMRI) of the rat brain. Visual stimulation at higher frequency than the flicker fusion frequency threshold leads to continuity illusion and is associated with negative BOLD response in the visual cortex, triggered by the superior colliculus. In this paper, we explored this mechanism using fMRI of the rat brain with visual stimulation at low (1Hz) and high (25Hz) frequency. We compared responses between different brain regions (the dorsolateral geniculate nucleus of the thalamus, the medial and lateral parts of the superior colliculus, and the visual cortex), sexes, and field strengths (9.4T and 14T, with varying contributions from large vessels). Results confirmed distinct neural responses to low and high frequency stimulation and highlighted the role of the lateral part of the superior colliculus in the transition from static to dynamic vision. Finally, we evaluated the ability of apparent diffusion coefficient (ADC)-fMRI to detect response to visual stimulation without vascular contribution. We found significant ADC-fMRI response in the medial and lateral parts of the superior colliculus but also in the corpus callosum. Our results highlight the ADC-fMRI high spatial specificity and high sensitivity to white matter.

## 1 Introduction

The visual system of the human brain exhibits remarkable plasticity, enabling adaptive responses to various environments and stimuli. As in other mammals, such as rodents, visual perception is assured by the pathway starting from the retinal neurons projecting to the brain visual areas^1^. The decoding and extraction of visual features for high-order tasks such as motion detection or object recognition is established by the cross-talks between regions of subcortical and cortical grey matter constituting the visual pathway^2,3^. Retinal neurons project mainly to the dorsolateral geniculate nucleus (DLGn) of the thalamus and to the superior colliculus (SC) which are both interconnected with the visual cortex^4,5^. While the functional architecture of these regions has been described^6^, it is still unclear how the interplay between these subregions drives the adaptive response to varying stimuli.

In particular, the mechanisms underlying the transition from perception of individual events to a dynamic vision, for instance of moving objects are still unknown. This transition can be described by the flicker fusion frequency (FFF) threshold which is defined, for a flashing stimulus, as the frequency at which the individual stimuli are perceived as one^7,8^. While electrophysiological studies have shown that the FFF threshold depends on the retinal organization of a species^9,10^, behavioural measurements of FFF have yielded very different values, suggesting a cortical contribution to the definition of the threshold^11^.

As an alternative technique, Blood Oxygen Level Dependent (BOLD) functional MRI (fMRI)^12^ can be used to study whole-brain activity in response to a task. The BOLD signal increases following vascular response to neuronal firing^13^. This neurovascular coupling involves complex changes in local blood flow, volume, and oxygenation aimed at supporting the metabolic demands of firing neurons^12,14,15^. Hence, BOLD-fMRI studies indirectly detect neuronal activity through its vascular response, with an intrinsic low spatial and temporal specificity^16^, as well as reduced sensitivity in less vascularized areas such as white matter^17,18^.

To bridge the gap between electrophysiological and behavioural FFF values, BOLD-fMRI studies have thus assessed the activation in the rat brain in response to a visual stimulus^19,20^. The spatial activation maps were reported to vary with increasing stimulation frequency. Starting from a positive BOLD response in both the visual cortex and the SC at low frequencies, as the frequency was increased and the FFF threshold was reached, the BOLD response became negative in the visual cortex. Further increasing the frequency resulted in a negative BOLD response in the SC as well. While positive BOLD response is a proxy for neuronal activity, negative BOLD can have multiple origins such as suppression or passive reduction of blood oxygenation in areas not relevant to the task, with less defined specificity to neural activity^21–23^.

Apparent diffusion coefficient (ADC)-fMRI^24,25^ has been proposed as an alternative contrast to probe neuronal activity more directly than its BOLD counterpart by recording changes in the diffusion properties of the tissue resulting from neuromorphological coupling, i.e. microstructure fluctuations concomitant with neuronal activity . Recent advances in the acquisition schemes have allowed the minimization of haemodynamic contributions to the ADC-fMRI timecourses^29–31^. In particular, the computation of ADC from two diffusion-weighted spin echo fMRI time series with alternating b-values substantially reduces T_2_ contributions to the signal, the use of b≥200 s/mm^2^ suppresses direct blood contributions, and the use of cross-term compensated gradient waveforms minimizes contributions from background blood susceptibility gradients^30,31^. In addition to minimizing vascular confounds, ADC-fMRI has recently demonstrated the ability to detect neuronal activation in the human brain in response to motor and visual stimulations with enhanced sensitivity to white matter^30^. However, in the mentioned study, a linear diffusion encoding was used, leading to spatial dependence of ADC-fMRI sensitivity and favouring white matter tracts that were perpendicular to the diffusion weighting direction^32,33^. To avoid fiber-orientation bias, spherical tensor encoding can be used to achieve isotropic diffusion encoding in one acquisition^33,34^.

In the present work, we characterized the functional interplay underlying FFF threshold in the rat brain using fMRI with visual stimulation at two frequencies: a low frequency of 1Hz at which the visual pathway operates in static vision mode (every flash is encoded as a separate event) and a high frequency of 25Hz reported to induce a shift to dynamic vision mode (flashing stimuli are fused together)^20^. For both stimulation frequencies, we compared BOLD and ADC-fMRI activation maps derived from diffusion-weighted images acquired using isotropic encoding. Neural responses were analysed both jointly and separately for males and females, to provide a comprehensive description of the neuronal suppression occurring at high frequency, engaging both grey and white matter areas. Additionally, we compared BOLD and ADC-fMRI activation maps acquired at 9.4T and 14T to explore the effect of higher blood magnetic susceptibility changes on each functional contrast with increasing field strength.

## 2 Materials and Methods

### 2.1 Animal experiments

#### 2.1.1 Animals

All experiments were conducted in respect of the Swiss law for protection of animals (OPAn) and were authorized by cantonal (authorization n° VD3958B) and national (Authorization n° 36256) ethics committees.

Sprague Dawley rats (n=18, 12 females and 6 males, 200-225g, 8-10 weeks old) were received from Charles River Laboratories. Rats were housed in cages of 2 (males) or 3 (females) with a 12h day/night cycle, water and food ad libitum, and nesting enrichment. All experiments were performed after a minimum of 1 week of acclimatization. Animals were split into three groups. The first and second groups consisted in 6 females and 6 males respectively and were scanned on a 14T Bruker MRI system. The third group, constituted of 6 females, was scanned on a 9.4T Bruker MRI system. The number of 6 animals per group was decided based on other studies with similar sample size^35^. No exclusion or inclusion criterion was used, data from all animals were included.

#### 2.1.2 Anaesthesia and monitoring protocol

Rats were initially anesthetized with isoflurane at 4% in a 70/30% air-oxygen mixture. Isoflurane was reduced to 2% after induction. Rats were then transferred and fixed using a bitebar and earbars to a homemade MRI cradle. Eye ointment was applied. Rats were shaved on the back and a catheter (22G) was inserted subcutaneously for medetomidine administration. Body temperature was continuously monitored using a rectal thermometer and maintained around (37 ± 0.5) °C using a warm water circulation system. Respiratory rate was also continuously monitored using a respiration pillow placed under the animal’s thorax and connected to a monitoring system. After installation of the rats in the MRI system, the anaesthesia was switched from isoflurane to medetomidine. First, a bolus of medetomidine (0.1 mg/kg) was injected via the subcutaneous catheter connected to a perfusion line and isoflurane was reduced to 0% gradually during the following 10 minutes^36–38^ aiming for a respiration rate ∼80-85 bpm. Fifteen minutes after bolus administration, a continuous subcutaneous infusion of medetomidine (0.1 mg/kg/h) was started and maintained until the end of the MRI experiment^39^. Maximum duration of the medetomidine anaesthesia was 3h. At the end of the last scan, the medetomidine perfusion was switched off and an intramuscular injection of atipamezole (0.5 mg/kg) was performed as antagonist to medetomidine before returning the rat to its cage.

### 2.2 Visual stimulation

Visual stimulation setup was similar for both 9.4T and 14T MRI scanners. A bifurcated optic fiber connected to a blue LED (λ=470nm and I=8.1×10−1 W/m^2^, Doric Lenses, Quebec, Canada) was placed horizontally in front of each eye of the animal in the MRI bed for binocular visual stimulation during the MRI experiment. The LED was piloted by an Arduino connected to the MRI output trigger. fMRI sequences triggered the following visual stimulation paradigm: after an initial 24s of rest, epochs of 16s of bilateral flashing light and 24s of rest were repeated 12 times for a total duration of 8m24s. The Arduino was resynchronized with the MRI trigger after each epoch to avoid accumulation of delay between the MRI and the internal Arduino clock. Visual stimulus was flashing at a frequency of either 1Hz or 25Hz, with a flash duration of 10ms^20^.

### 2.3 MRI experiments

#### 2.3.1 14T MRI

The first and second groups were scanned on a 14T Bruker MRI system (G_max_=1000 mT/m, S_max_=5600T/m/s) operating on ParaVision 360 v1.1 software (Bruker, Ettlingen, Germany) and using a volume coil for transmission and a 2-channel surface coil for reception (Rapid Biomedical GmbH, Rimpar, Germany). After animal installation and switch of anaesthesia, a B_0_ field map was acquired and used for shimming in an ellipsoidal volume covering the rat brain using MapShim. A T_2_-weighted (T_2_-w) image was acquired for anatomical reference (TurboRARE, TR/TE=2500/6ms, Resolution 0.125x0.125x0.5 mm^3^, Matrix 160x160, 45 slices, duration=6min40s). fMRI acquisitions started at least 30 min after the switch of anaesthesia from isoflurane to medetomidine to allow for isoflurane clearance. BOLD-fMRI was acquired using a gradient echo planar imaging sequence (GRE-EPI, TR/TE = 1000/11 ms, Resolution 0.38x0.38x1.5 mm^3^, Matrix 68x68, 10 slices, 504 repetitions) and ADC-fMRI using a diffusion-weighted spin echo EPI sequence (dw-SE-EPI, TR/TE=1000/41 ms, matched geometry with GRE-EPI, b_1_=200 s/mm², b_2_=1000 s/mm², 504 repetitions). Diffusion-weighting was performed using symmetric spherical tensor encoding with cross-term compensated waveforms. Eight fMRI runs were acquired in total for each rat: BOLD-fMRI and ADC-fMRI sequences were acquired twice for each stimulation frequency, in a randomized order. Twelve dummy repetitions were acquired and discarded in the beginning of every scan to reach steady state. Forward and reverse phase encoded images were also acquired for GRE-EPI and for dw-SE-EPI with a null b-value (b_0_ image) to enable EPI spatial distortion correction during pre-processing.

#### 2.3.2 9.4T MRI

The third group was scanned on a 9.4T Bruker MRI system (G_max_=660 mT/m, S_max_=4570 T/m/s) operating on Paravision 360 v3.5 software, using a volume coil for transmission and a 6-channel surface cryoprobe for reception. After installation, the shimming procedure was identical to that on the 14T system. Acquisition parameters for the anatomical image, GRE-EPI and dw-SE-EPI were kept as consistent as possible to the 14T protocol. Some variations were necessary to accommodate hardware differences between the two systems. Anatomical reference (TurboRARE, Resolution 0.14x0.14x1.1 mm^3^, 256x256, 20 slices, duration=10m40s), GRE-EPI (TR/TE = 1000/10 ms, Resolution 0.36x0.36x1.5, Matrix 69x50 with saturation band, 8 slices), and dw-SE-EPI (TR/TE = 1000/42 ms, matched geometry with GRE-EPI) were acquired using the same protocol. The same gradient waveforms as previously were used for diffusion-weighting with amplitude rescaling to adapt to the 9.4T gradient limitations.

**Table 1:** Sequence parameters used for anatomical reference and functional MRI, at 9.4T and 14T.

|  | Anatomical Reference |  | BOLD-fMRI |  | ADC-fMRI |  |
| --- | --- | --- | --- | --- | --- | --- |
|  | 9.4T | 14T | 9.4T | 14T | 9.4T | 14T |
| Sequence | TurboRARE |  | GRE-EPI |  | dw-SE-EPI |  |
| TR/TE (ms) | 2500/8 | 2500/6 | 1000/10 | 1000/11 | 1000/42 | 1000/41 |
| Resolution (mm<sup>3</sup>) | $0.14 \times 0.14 \times 1.1$ | $0.125 \times 0.125 \times 0.5$ | $0.36 \times 0.36 \times 1.5$ | $0.38 \times 0.38 \times 1.5$ | $0.36 \times 0.36 \times 1.5$ | $0.38 \times 0.38 \times 1.5$ |
| Slices | 20 | 45 | 8 |  | 8 |  |
| b-values (s/mm<sup>2</sup>) | - |  | - |  | 200, 1000 |  |
| Acquisition time (s) | 640 | 400 | 504 |  | 504 |  |

### 2.4 Image pre-processing

Raw data extracted from Bruker Paravision software were converted to NIfTI using the Dicomifier toolbox (https://github.com/lamyj/dicomifier). Following this step, dw-SE-EPI time series were split per b-values, leading to two different time series (b200 and b1000) of 252 repetitions with 2s temporal resolution each. BOLD time series corresponded to GRE-EPI time series of 504 repetitions with 1s temporal resolution. BOLD, b200 and b1000 time series underwent the same pre-processing steps including MP-PCA denoising^42^, Gibbs unringing ^43^, distortion correction^44^, and motion correction (rigid + affine)^45^.

Following these steps, the pre-processed b1000 time series was registered to the b200 time series, and the resulting aligned datasets were used to compute the ADC time series. The brain was extracted from the T_2_-w image using RATS^46^ to yield a brain mask. The T_2_-w image was then registered to functional images (average of all BOLD volumes for GRE-EPI, average of all b200 repetitions for dw-SE-EPI) using ANTs^45^ and the transformation was applied to the brain mask for skull stripping of functional time series. High-pass filtering (100s cutoff) was applied to functional time series at this step. No spatial smoothing was performed.

Multivariate templates were generated from the T_2_-w, b_0_, and average BOLD images of the six rats from the second cohort^47^. Individual functional time series were registered to their corresponding (b_0_ or BOLD) template using ANTs^45^. Waxholm Space Atlas of the rat brain was first registered to the T_2_-w template and then propagated into subject space by applying the inverse transformations previously estimated in order to obtain for segmentation of the Optic tract (Ot), Corpus callosum (CC), Dorsolateral geniculate nucleus (DLGn), Superior colliculus (SC), Primary and secondary visual cortices (V1 and V2), and retrosplenial cortex (RSC). The Superior Colliculus was manually segmented on the Waxholm Space Atlas between its medial and lateral parts using ITK-snap (Supplementary Figure 1).

### 2.5 FMRI analysis

#### 2.5.1 BOLD activation maps

Processed BOLD time series underwent general linear model (GLM) analysis at the subject-level using a boxcar response function and cluster-correction with threshold of |Z|>2.3 and p<0.05. As the hemodynamic response in the rat brain is much faster than in humans, the boxcar response provides a better fit than traditional HRF^48^. Results from the subject-level GLM were then registered to the template space. A group-level GLM analysis was performed separately per field strength, stimulation frequency and, in the case of 14T data, per sex on the registered Z-score maps, using a cluster-correction of either |Z|>1.5 or |Z|>2.3 and p-value threshold of 0.05.

In order to make results more directly comparable to ADC-fMRI analysis (see below), BOLD timecourses from female rats at 9.4T also underwent GLM analysis using the Finite Impulse Response (FIR) method.

#### 2.5.2 ADC activation maps

To avoid assumptions on the ADC-fMRI response function, subject-level GLM analysis was performed on ADC time series with boxcar response function convolved with FIR (4 impulses, 16s window)^49^ and cluster-correction with threshold of |Z|>2.3. As FIR analysis does not yield positive or negative contrasts like boxcar, responses in significant voxels were pooled across rats for each field strength and each stimulation frequency and classified using time series K-means clustering with 10 clusters and k-means++ initialization. The average response was extracted per cluster. Clusters with average positive or negative values during visual stimulation were pooled together for better visualization of voxels with a positive or negative ADC response.

#### 2.5.3 Average response plots

To visualize the average response to visual stimulation in specific brain regions, for each time series the signal was first averaged across voxels included in the mask of the region, normalized to the mean value of the first 6 volumes of each epoch, and averaged across epochs. Finally, the mean value and standard deviation across time series were computed and visually represented.

For better visualization and comparison with BOLD, ADC time series with a temporal resolution of 1s were also generated. To that end, we first linearly interpolated the b200 and b1000 time series to a 1s resolution. A new ADC time series was then computed from pairs of one interpolated and one measured b-value corresponding to the same time-point in the series. The resulted interpolated ADC time series were only used for plotting the average ADC response. The GLM and FIR analyses described above were performed exclusively with the original 2s temporal resolution.

## 3 Results

### 3.1 14T BOLD-fMRI shows specific activation patterns in response to 1Hz and 25Hz visual stimuli

Activation of the visual system was probed in male and female rats at 14T using visual stimulation. Two different stimulation frequencies were used to investigate responses to static (1Hz) and dynamic (25Hz) visual stimuli. Figure 1 shows the group-level BOLD response to 1Hz and 25Hz visual stimulation in males, females, and in all animals combined. Visual stimulation with a frequency of 1Hz (Figure 1A-C) induced a large area of strong positive BOLD response centred on the DLGn and SC, as well as a positive BOLD response with lower intensity in V1 and V2. Activation maps of females (Figure 1A) and males (Figure 1B) were spatially consistent, however significant BOLD response covered a larger area in females than in males. Cortical areas with significant negative BOLD response were also found in a location corresponding to AuC.

**Figure 1:**
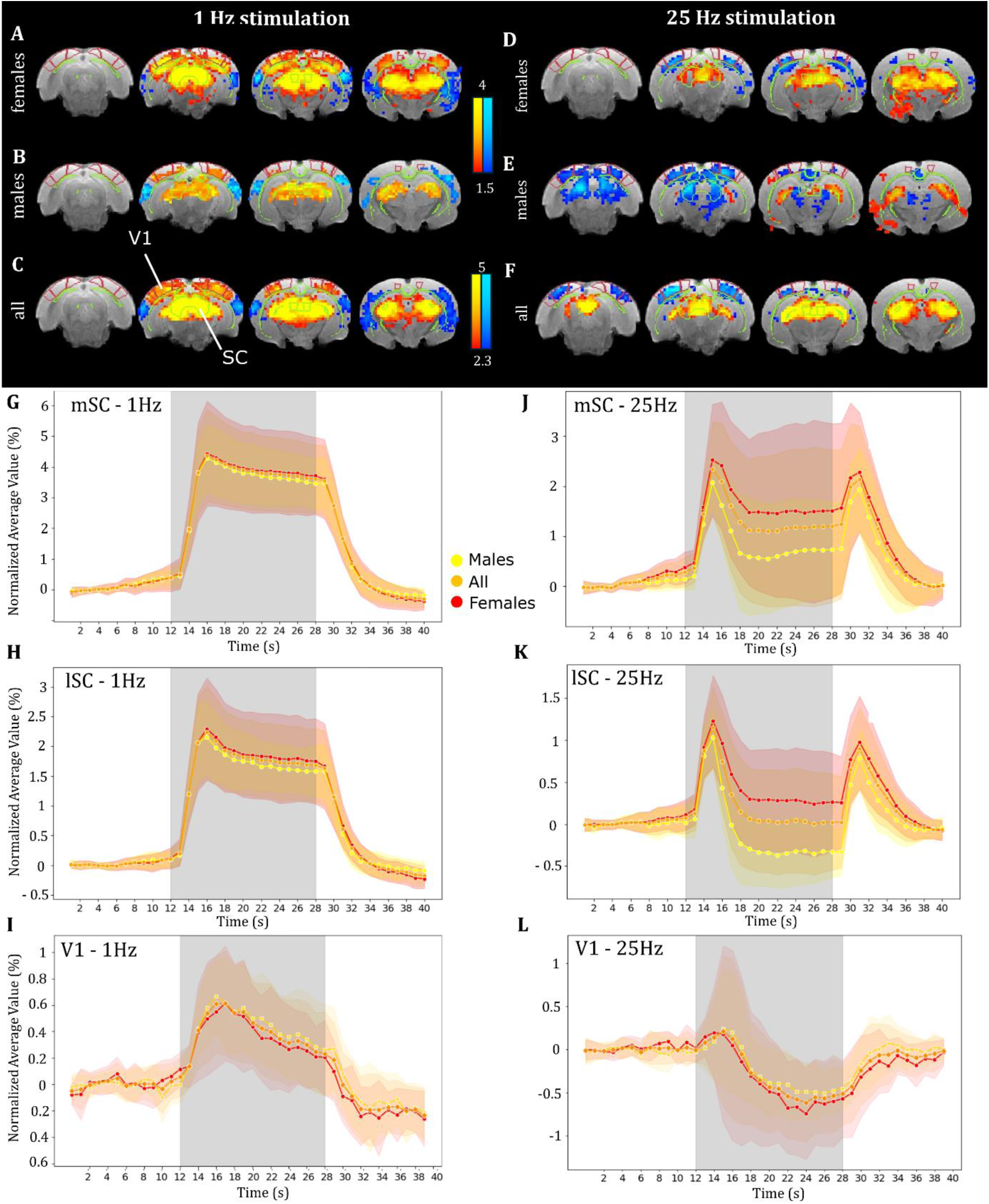
14T BOLD response to 1Hz and 25Hz visual stimulation in the brain of male and female rats. A-F: Cluster-corrected group-level activation maps of significant positive (red- yellow) and negative (blue-light blue) BOLD response to 1Hz (A-C) and 25Hz (D-F) visual stimulation in (A & D) females only (n=6, |Z|>1.5), (B & E) males only (n=6, |Z|>1.5), (C & F) all animals (n=12, |Z|>2.3). G-I: Average response to 1Hz visual stimulation in (G) the medial SC, (H) the lateral SC, and (I) V1 in females (red, n=6), males (yellow, n=6), and all animals (orange, n=12). J-L: Average response to 25Hz visual stimulation in (J) the medial SC, (K) the lateral SC, and (L) V1. Shading denotes the standard deviation. Grey area represents stimulus duration.

To visualize the activation with respect to stimulus timing, we plotted the average response to 1Hz stimulation in the SC and V1. The average BOLD response in the SC (Figure 1G-H) showed very similar temporal dynamics between males and females, differing primarily in response magnitude. The maximum amplitude of the BOLD response was higher in the medial part (4.5%, Figure 1G) than in the lateral part of the SC (2.5%, Figure 1H), in both sexes. The BOLD response in V1 (Figure 1I) was less intense in comparison to SC. The maximum amplitude change was 0.6% in both sexes and the plateau effect was less visible. After the end of the stimulus the amplitude decreased and reached negative values compared to baseline.

In response to 25Hz visual stimulation, the visual cortex presented a negative BOLD response, contrasting with 1Hz stimulation. This observation was spatially consistent in females (Figure 1D) and males (Figure 1E). In contrast, in the SC, activation maps showed a positive response in females and a negative response in males. In females, the positive response was constrained to the medial SC while in males the negative response was constrained to the lateral SC. When taking both males and females into account, activation maps showed areas of positive BOLD response in the medial part of the SC only and in the DLGn (Figure 1F). The spatial extent of the activation in these areas was narrower than with the 1Hz stimulation.

Average response to the 25Hz stimulus in the SC (Figure 1J-K) revealed a disparity between males and females. The global shape of the response was consistent in both sexes, with two peaks at the start and at the end of the stimulus, separated by a plateau. Again, the amplitude of the response was higher in the medial SC (Figure 1J) than in the lateral SC (Figure 1K). In both subregions, the maximum amplitude change was higher in females than in males. In males, the response in the lateral SC plateaued below the baseline, which explains the negative activation found in the lateral part of the SC (Figure 1F). Finally, in V1, the response to 25Hz stimulation (Figure 1J) showed an initial low amplitude (0.2%) positive response during the first 4s of the stimulus, followed by a sustained decrease into negative BOLD until the end of the stimulus. Again, this effect was stronger in females (-0.6%) than in males (-0.4%).

### 3.2 BOLD response to visual stimulation shows distinct responses despite consistent activation maps at 9.4T vs 14T

The visual stimulation paradigm was reproduced in female rats on a 9.4T MRI to test the robustness of the activation maps found at 14T and to study the influence of the field strength on the BOLD response to both 1Hz and 25Hz stimulations (Figure 2).

**Figure 2:**
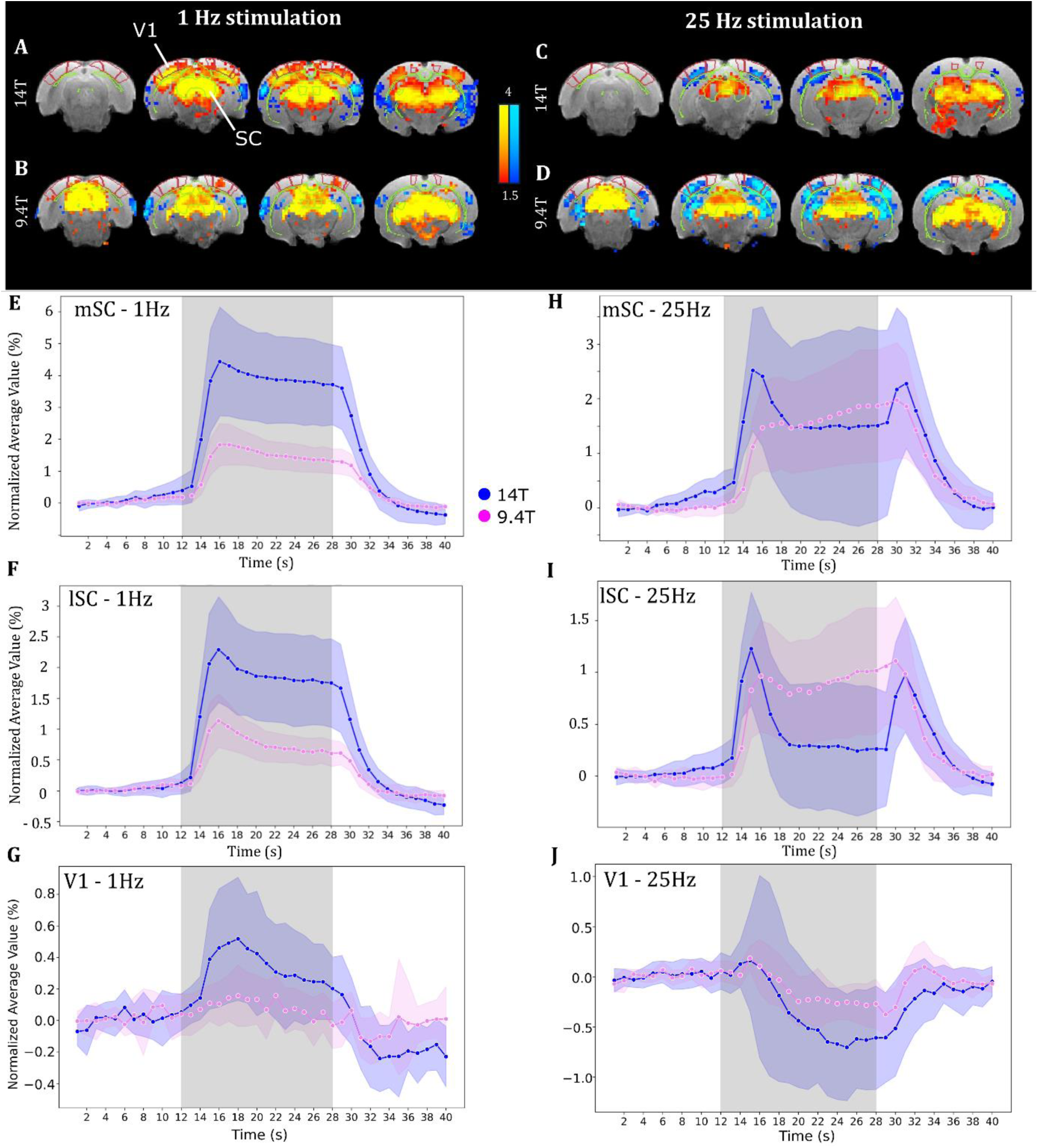
BOLD response to 1Hz and 25Hz visual stimulation in female rats at 9.4T and 14T. A-D: Cluster-corrected group-level activation maps of significant positive (red-yellow) and negative (blue-light blue) BOLD response to 1Hz (A-B) and 25Hz (C-D) visual stimulation at (A & C) 14T (n=6 females, |Z|>1.5), (B & D) 9.4T (n=6 females, |Z|>1.5). E-G: Average response to 1Hz visual stimulation in (E) the medial SC, (F) the lateral SC, and (G) V1 at 14T (blue, n=6) and 9.4T (magenta, n=6). H-J: Average response to 25Hz visual stimulation in (H) the medial SC, (I) the lateral SC, and (J) V1. Shading denotes standard deviation. Grey area represents stimulus duration.

In agreement with the activation map obtained from 14T MRI (Figure 2A), visual stimulation with a frequency of 1Hz induced a large area of positive BOLD response centred on the SC and DLGn at 9.4T (Figure 2B). In contrast, the area with significant positive BOLD response detected in the visual cortex at 14T was reduced to a unilateral narrow cluster in right V1 at 9.4T. Negative BOLD was also detected in the auditory cortex and was consistent across field strengths.

The average response to 1Hz stimulation in the SC (Figure 2E-F) was similar in shape across field strengths. The maximum amplitude of the response was reduced at 9.4T compared to 14T (2% vs 4.5% in the medial SC, 1.2% vs 2.5% in the lateral SC). The average response in V1 (Figure 2G) shows a very mild BOLD response at 9.4T with a maximum amplitude of only 0.2% compared to 0.6% at 14T, which may explain the smaller area with significant BOLD response on the activation map.

The specific activation pattern obtained in females at 14T with 25Hz visual stimulation (Figure 2C) was mostly consistent at 9.4T (Figure 2D). An area of positive BOLD response was found in the SC and DLGn but seemed to be centred on the lateral part of the SC, while at 14T it was mostly the medial part of the SC which showed significant activation. In addition, negative BOLD was found in a much larger proportion at 9.4T than at 14T in cortical areas encompassing V1 and V2, and in subcortical areas such as the hippocampus.

Investigating the average response to 25Hz stimulation in the SC (Figure 2H-I) unveiled a marked difference in the shape of the response between the two field strengths. At 9.4T, the response did not display a double peak pattern but instead presented an initial large increase, followed by slower continuous increase in amplitude during the stimulation. For the same field, the response was similar between the medial and lateral SC.

In V1, the response to 25Hz stimulus was comparable between 9.4T and 14T, with the only difference being a reduced amplitude at 9.4T (Figure 2J).

### 3.3 ADC-fMRI detects specific signatures of visual stimulation in both subcortical grey matter and white matter at 9.4T

We investigated the ability of ADC-fMRI to probe response to a visual stimulus independently from the BOLD effect at 9.4T for both stimulation frequencies. We used a FIR function followed by unsupervised clustering to identify and isolate the voxels showing a significant signal variation during stimulation with low prior on the shape of the response function (Figure 3).

**Figure 3:**
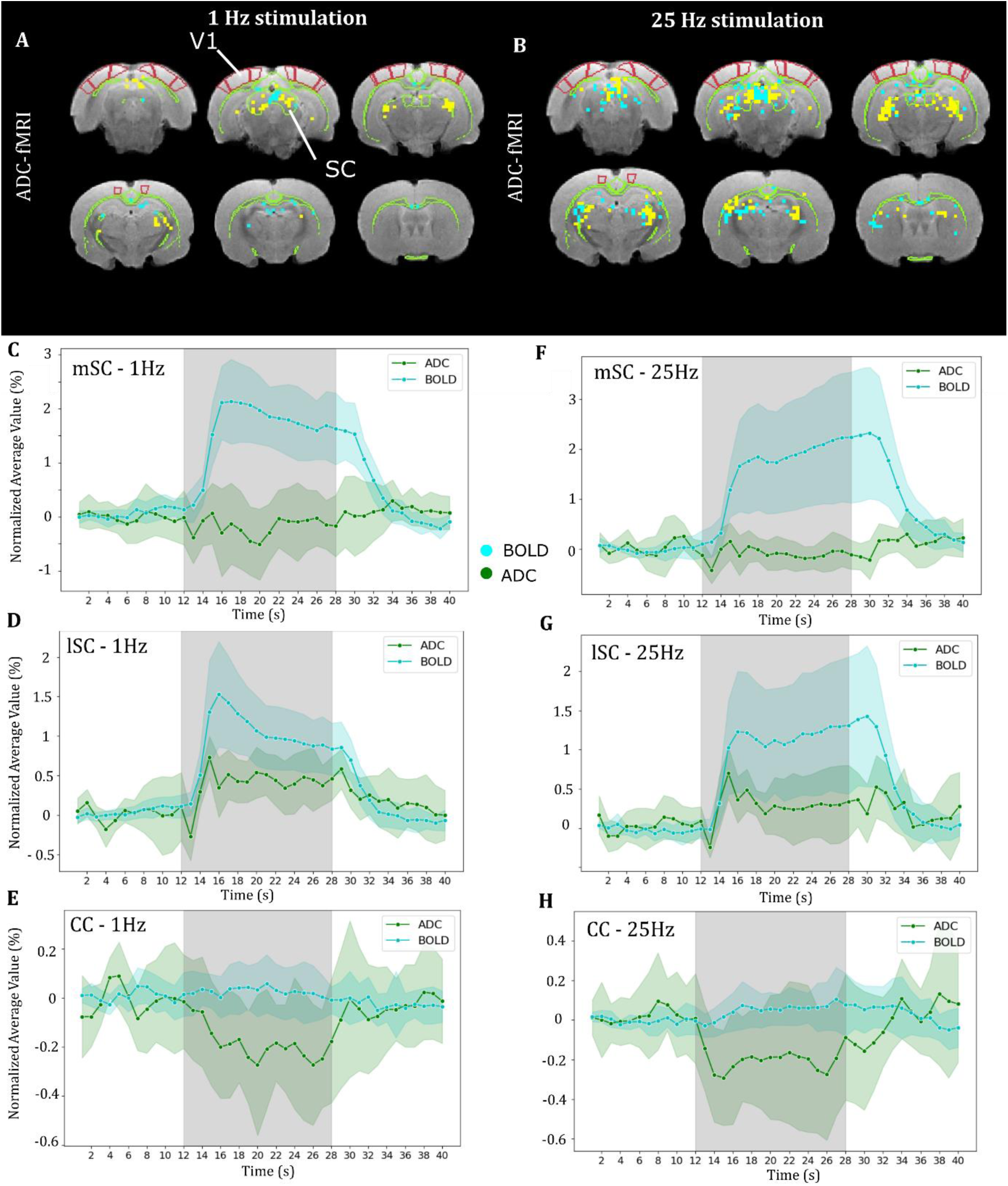
ADC response to 1Hz and 25Hz visual stimulation in comparison to BOLD at 9.4T. A- B: K-means clustered voxels with average response to (A) 1Hz and (B) 25Hz visual C-E & F-H: Average response to (C-E) 1Hz and (F-H) 25Hz visual stimulation in (C & F) voxels with negative ADC response detected in the medial SC, (D & G) voxels with positive ADC response detected in the lateral SC, (E & H) voxels with negative ADC response detected in the corpus callosum (CC). Shades represent standard deviation. Grey area represents stimulus duration.

In response to the 1Hz visual stimulus, clusters with negative ADC response were found in the medial part of the SC, the RSC and the CC, while positive ADC response was found in the lateral part of the SC and in the DLGn (Figure 3A). Comparing ADC and BOLD average response in these group-level clusters, we found negative ADC-fMRI in the medial SC in the presence of positive BOLD response (Figure 3C). The maximum amplitude of the ADC response was −0.5%, while the maximum amplitude of the BOLD response in the same area was 2.2%. Similar clusters were also found in the hippocampus. In the lateral part of the SC and in the DLGn, the positive BOLD response (+1.5% maximum amplitude) was accompanied by a positive ADC response with a very similar shape to BOLD and a maximum amplitude of +0.7% (Figure 3D). Remarkably, a negative ADC response (-0.3% maximum amplitude) was also detected in some clusters in the CC (Figure 3E), whereas no BOLD response could be observed.

Interestingly, ADC-fMRI yielded very similar spatial patterns between the 25Hz and 1Hz visual stimulation (Figure 3B). Clusters with negative ADC response to 25Hz stimulation were found in the medial part of the SC and in the hippocampus. These voxels covered a larger area as compared to the 1Hz stimulus, but showed lower amplitude (Figure 3F, −0.2%, vs −0.4% at 1Hz). In the same voxels, BOLD-fMRI showed a positive response (+2.2% maximum amplitude). Similarly to the 1Hz stimulus, the average ADC response in the lateral part of the SC was positive and showed a very similar shape to the BOLD response, with a maximum amplitude of +0.6% and +1.2% respectively (Figure 3G). Finally, negative ADC response was also found in clusters in the corpus callosum with a maximum amplitude of −0.2% in the absence of any BOLD response (Figure 3H).

The activation patterns highlighted by ADC-fMRI at 9.4T were unique to this functional contrast. Indeed, an identical analysis pipeline of BOLD-fMRI at 9.4T using FIR yielded similar maps to the boxcar GLM (Supplementary Figure 2).

However, activation patterns highlighted by ADC-fMRI at 14T resembled more closely those of BOLD, with the medial SC also showing a positive ADC-fMRI response at 14T, suggesting a more marked BOLD contamination at higher field strength (Supplementary Figure 3).

## 4 Discussion

The neural mechanisms underlying flicker fusion frequency are debated. Functional MRI has the potential to bring understanding of the phenomena at the network scale, thus bridging the gap between electrophysiological and behavioural measurements. However, the BOLD-fMRI contrast is driven by vascular response as an indirect measurement of neural response. In the present work, we explored the variations of BOLD-fMRI signature of the FFF threshold across sex and field strengths and found substantial differences that arise from the vascular origin of BOLD-fMRI. Interestingly, these differences highlighted a different response between the medial and lateral parts of the SC. Using ADC-fMRI as a more direct measurement of neural firing, we were able to detect activation in the visual network and in white matter. Moreover, ADC-fMRI also discriminated the medial and lateral parts of the SC.

### 4.1 14T BOLD-fMRI uncovers sex- and frequency-dependent response reflecting neuronal activation and suppression in the visual pathway

Using 14T BOLD-fMRI, we found overwhelming activation in the main brain structures of the rat visual pathway (DLGn, SC, V1) in response to visual stimulation with a frequency of 1Hz. In addition to this large positive BOLD response, we found a negative response in the auditory cortex. This phenomenon may arise from a suppression of the activity in regions that are not relevant to the task^23^. Signals from both sensory subcortical^50^ and high-order cortical^21^ areas have been identified as triggers of this negative response.

As previously observed^20,35^, the increase in stimulation frequency induced a transition to negative BOLD in the visual cortex. However, in our study, the BOLD response to 25Hz visual stimulation exhibited greater complexity than previously reported^20^ , especially in the SC. Our results showed for the first time that the response in the SC differed between males and females. While the BOLD response was still positive throughout the SC in females, a negative BOLD response was visible only in the lateral part of the SC in males at 25Hz. We found this difference to be mainly due to variations in the amplitude of the plateau between the two positive BOLD peaks that systematically occurred right after onset and end of the stimulus and solely at this frequency.

Medial and lateral parts of the SC show different organizations and integrate separately to specific subnetworks^51,52^. The medial part of the SC receives most of the input from V1 as part of the visual integration subnetwork, while the lateral SC is more integrated in visuomotor, sensorimotor, and escape-approach subnetworks. Our results highlight the different behaviour of these two subregions in response to the 25Hz stimulus, and suggest a more pronounced suppressing effect in the lateral SC than in the medial SC.

Interestingly, compared to Gil et al. 2024^20^, and despite using a similar experimental paradigm and stimulation frequencies, we observed residual positive BOLD response in the SC at 25 Hz. This difference suggests that Sprague-Dawley rats present a higher FFF threshold than Long-Evans rats, possibly due to their albino condition. However, this finding could also be attributed to the younger average age of the rats in our study.

### 4.2 BOLD dependency on B_0_ field strength highlights distinct response to 25Hz stimulus in subregions of the SC

Our results showed robust detection of both positive and negative BOLD areas across the investigated field strengths. However, one feature specific to the 14T BOLD response at 25Hz stimulation was the double-peak shape after onset and at the end of the stimulus in the SC. At 9.4T, this response showed a very different shape with monotonous increase at onset and decrease at offset.

Relative contributions from the microvascular, macrovascular, and extravascular compartments to the BOLD signal are highly dependent on the B_0_ field strength^53^. Extravascular contribution to the BOLD signal is dominant at ultra-high-field due to an increased extent of field distortions perpendicular to large radius vessels^14,54^. Thus, discrepancies between responses at 9.4T and 14T may reflect specific involvements from areas close to large vessels. Vascularization of the subcortex in this area consists in large vessels running dorsal to the SC and DLGn and supplying in depth the nuclei through small penetrating vessels^19,55^. Using monocrystalline iron oxide nanoparticles injection for MR angiography of the rat brain, Lau et al. showed that large vessels are closer to the DLGn and lateral SC than to the medial part of the SC^19^. This vascularization leads to an increased sensitivity to extravascular compartment in the lateral SC at 14T. Thus, it seems that the negative BOLD response observed in Figure 1K originates from the extravascular compartment in the vicinity of the large vessels running near the lateral SC. This result is also consistent with the differences in activation maps between sexes, showing negative BOLD in the lateral SC in males only (Figure 1F). Cerebral arteries show larger inner diameter in male rats than in females^56^, while cerebral blood volume and vascular density are higher in females^57^. These differences in vasculature lead to distinct BOLD responses between males and females, which can be a confounding factor for the interpretation of BOLD activation in terms of neuronal firing.

### 4.3 ADC-fMRI detects neuronal activation with high spatial specificity and sensitivity to white matter at 9.4T

Our FIR analysis of ADC-fMRI was able to detect voxels responding to the task with reduced assumption on the shape of the response. The identified voxels were mostly localized in the visual pathway, suggesting that despite high freedom in the response shape, this analysis was still specifically detecting voxels responding to the task.

In order to visualize this response, we applied a clustering algorithm to the time series of identified voxels. Strikingly, the resulting activation maps at 9.4T were very similar between 1Hz and 25Hz stimulation. This further reduces the probability for these significant areas to be false positives. Negative ADC response was found in the medial part of the SC at both frequencies with different amplitudes. An ADC decrease has been previously shown to result from excitatory neuronal activity, even in the absence of vasculature^32,58–61^. This is corroborated by the strong positive BOLD response observed in the same brain area.

A negative ADC response was also found in the hippocampus. The hippocampus, while not considered part of the visual pathway per se, is involved in high-order visual processes^62^. In particular, in the rodent brain, the hippocampus is connected to the medial part of the SC as part of the head direction network^51^. While these voxels are packed within the large area of positive response in the SC using BOLD-fMRI, in ADC-fMRI they constitute a distinct cluster. This result highlights the potential of ADC-fMRI to detect activation with improved spatial specificity^30^. Of note, a previous study of rat forepaw stimulation reported rather an increase in ADC in the subcortical regions involved, directly or indirectly, in the somatosensory pathway – thalamus, hippocampus, striatum – in the absence of a detectable BOLD response in those areas^63^. The polarity of the ADC response may yield an accurate indication of tuneable excitatory/inhibitory balance in subcortical regions participating in a variety of sensory and higher-order function pathways^59,60^.

Negative ADC response was also detected in a few clusters located in white matter, and more specifically in the corpus callosum and hippocampal commissure, suggesting neuronal firing of interhemispheric projections which may be linked to bilateral stimulation. In comparison, no specific activation of white matter was detected using BOLD-fMRI at either field strength and using either boxcar or FIR response functions. This is consistent with similar recent findings of ADC-fMRI activation in the human optic radiation^33^. While the association between the detected white matter bundles and the task needs more investigation, this result reinforces previous findings from both task and resting-state fMRI^29,30^, highlighting the improved sensitivity of ADC-fMRI to white matter neural response.

Overall, our results demonstrate, for the first time, the detection of activation in rat brain regions using ADC-fMRI contrast, without a BOLD counterpart. In agreement with previous studies, we show that ADC-fMRI acquisition designs that minimize vascular contributions lead to improved spatial specificity and sensitivity to white matter activity.

### 4.4 Limitations and perspectives

Arterial dilation occurring at the beginning of the hemodynamic response could lead to a reduction in ADC due to increased partial volume effect from blood vessels inside a voxel. While a contribution of this effect to ADC-fMRI detection cannot be totally ruled out, it is very unlikely that it is the main phenomenon detected here, as ADC-fMRI response was detected solely in regions with reduced vascularization.

In addition to the negative clusters, positive ADC was found in the lateral SC and in the DLGn at 9.4T. The shape of the ADC response was very similar to the BOLD response in these regions for both stimulation frequencies. Such similarity strongly implies a BOLD contribution to the ADC time series, despite our efforts to minimize them. Interestingly, a negative ADC response was nonetheless found in the medial SC, where the positive BOLD response was the strongest. As discussed above, the vascularization of the lateral and medial parts of the SC are very different. This suggests that vascular contribution to ADC response highly depends on the proximity to large vessels. Furthermore, at 14T, where the BOLD effect is even more pronounced, ADC response maps largely mirrored BOLD maps, suggesting more widespread BOLD contamination.

Reducing BOLD contribution is crucial to fully leverage the higher specificity of ADC-fMRI. In this study, the BOLD contribution to ADC was minimized using cross-term compensated waveforms, calculating an ADC from interleaved b-values, instead of single b-value diffusion-weighted (and T_2_ weighted) timecourses, and employing a minimal b-value of 200 s/mm² to reduce direct signal from the intravascular compartment. However, several measures may still be taken for further minimizing the BOLD contribution. Firstly, the acquisition of alternative b-values leads to imperfect T_2_-weighting cancellation in the computation of the ADC, as the T_2_ may vary between two successive images used to compute one ADC volume. In order to accurately cancel the T_2_- weighting, the acquisition needs to be modified to either acquire both b-values within the same TR^64^ or to acquire each b-value with two echoes to estimate and correct for the T_2_-weighting. Secondly, as already mentioned, as the amplitude of the BOLD signal heavily relies on field strength, it is expected that reducing the field strength will lead to a reduction of this contribution. Indeed, we observed higher BOLD contributions to ADC-fMRI at 14T than at 9.4T. In comparison, previous literature studies on 3T ADC-fMRI were relatively insensitive to BOLD^29,30,65^. However, higher field strength offers increased sensitivity, which is particularly advantageous for ADC-fMRI given its inherently low response amplitude.

To conclude, in this study we investigated the neuronal response to visual stimuli in the rat using different frequencies, field strengths, and functional contrasts for comprehensive characterization. For the first time, we demonstrated the ability of ADC-fMRI to map the neural response to a visual task in the rat brain, and particularly in white matter where the response was not matched by the BOLD contrast. Based on the BOLD- and ADC-fMRI response dependency to field strength and sex, we were able to identify subregions of the SC with distinct vascularization and functional integration. Further studies are required to confirm our results suggesting that inhibition of the SC observed at stimulation frequency higher than the FFF threshold originates from the lateral part of the SC. Additionally, our results highlighted the potential confounding effect of vasculature on BOLD contrast, and the need to further reduce BOLD contribution to ADC-fMRI at ultra-high field in highly vascularized brain areas, in order to fully exploit the sensitivity and specificity of ADC-fMRI under optimal conditions.

## Acknowledgments

We acknowledge the CIBM Center for Biomedical Imaging co-founded and supported by Lausanne University Hospital (CHUV), University of Lausanne (UNIL), École polytechnique fédérale de Lausanne (EPFL), University of Geneva (UNIGE) and Geneva University Hospitals (HUG) for providing expertise and resources to conduct this study. In particular, we would like to thank Cristina Cudalbu, Thanh Phong Lê, Estelle Gerossier and Stefan Mitrea. We would also like to thank Rita Gil, and Francesca Barcellini for advice on the visual stimulation and Arduino interfacing. This work was supported by ERC Starting Grant ‘FIREPATH’, SERI no. MB22.00032. IJ is supported by Swiss National Science Foundation (SNSF) Eccellenza fellowship no. 194260. V.Z. and N.M are supported by SNSF Eccellenza PCEFP3_203005.

## Author Contribution

J-B.P.: conceptualization, data curation, formal analysis, investigation, methodology, visualization, and writing original draft; A.H.: Data curation, methodology, investigation, writing - review and editing; A.S., J.N.-D., N.M., V.Z, M.Y.: methodology, writing - review & editing; I.J.: conceptualization, funding acquisition, project administration, resources, supervision, and writing original draft.

## Data availability

The data, code, protocols, and key lab materials used and generated in this study are listed in a Key Resource Table alongside their persistent identifiers at https://doi.org/10.5281/zenodo.15705581.

## SUPPLEMENTARY FIGURES

**Supplementary Figure 1:**
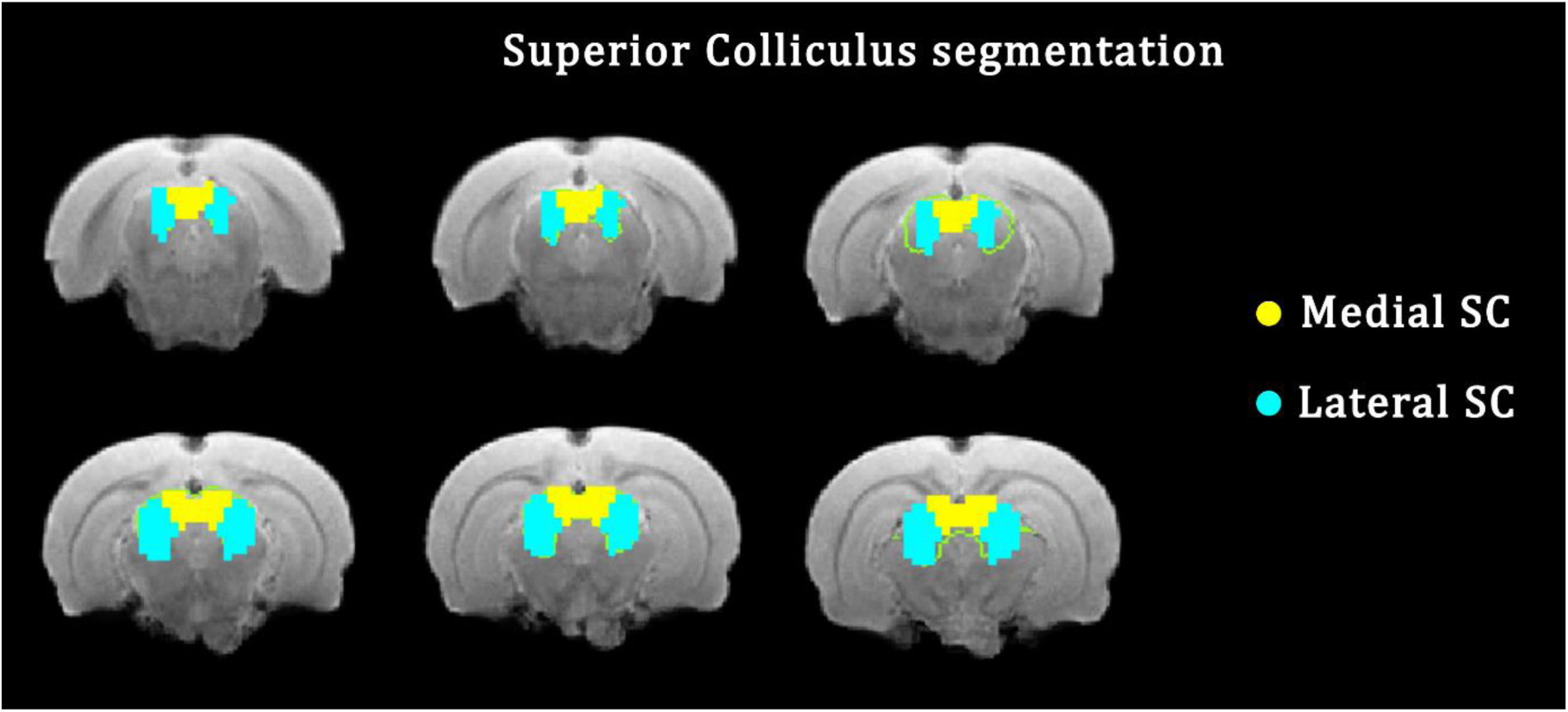
Segmentation of the medial and lateral parts of the superior colliculus (SC)

**Supplementary Figure 2:**
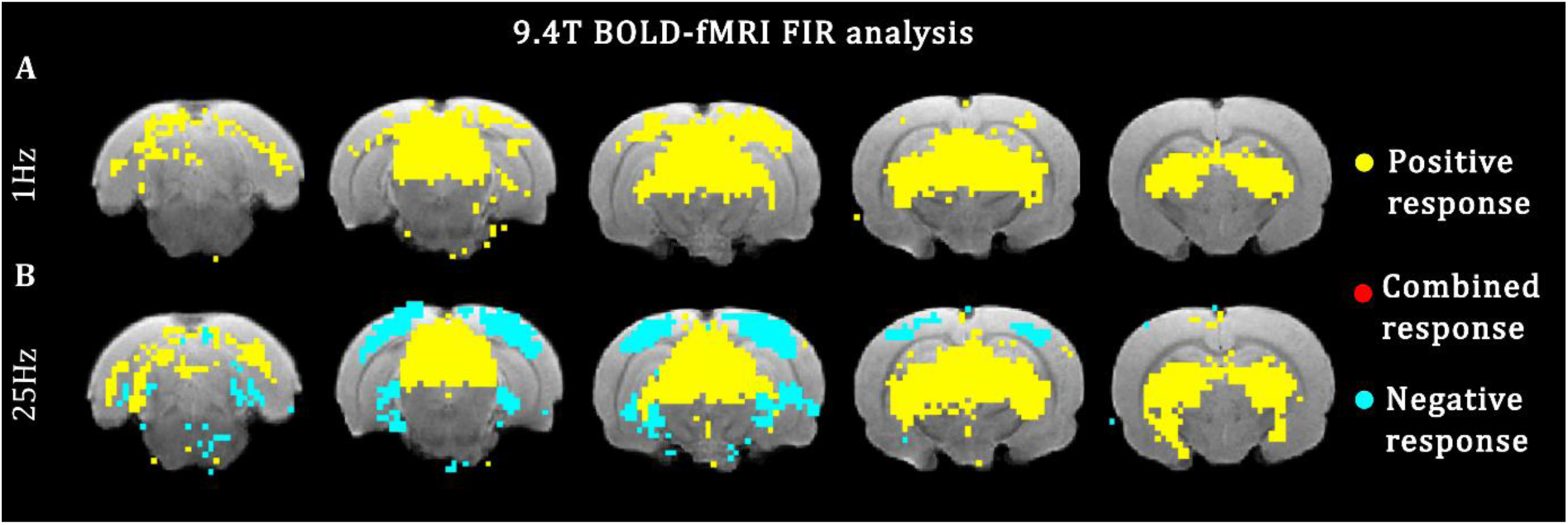
Activation maps of BOLD-fMRI resulting from the GLM analysis using a FIR response function at 9.4T in female rats, in response to (A) 1Hz and (B) 25Hz visual stimulation, mirroring the ADC-fMRI FIR analysis (Figure 3A-B). Similar information to GLM with a boxcar response function is found (Figure 2B-D). No specific activation in white matter regions can be observed.

**Supplementary Figure 3:**
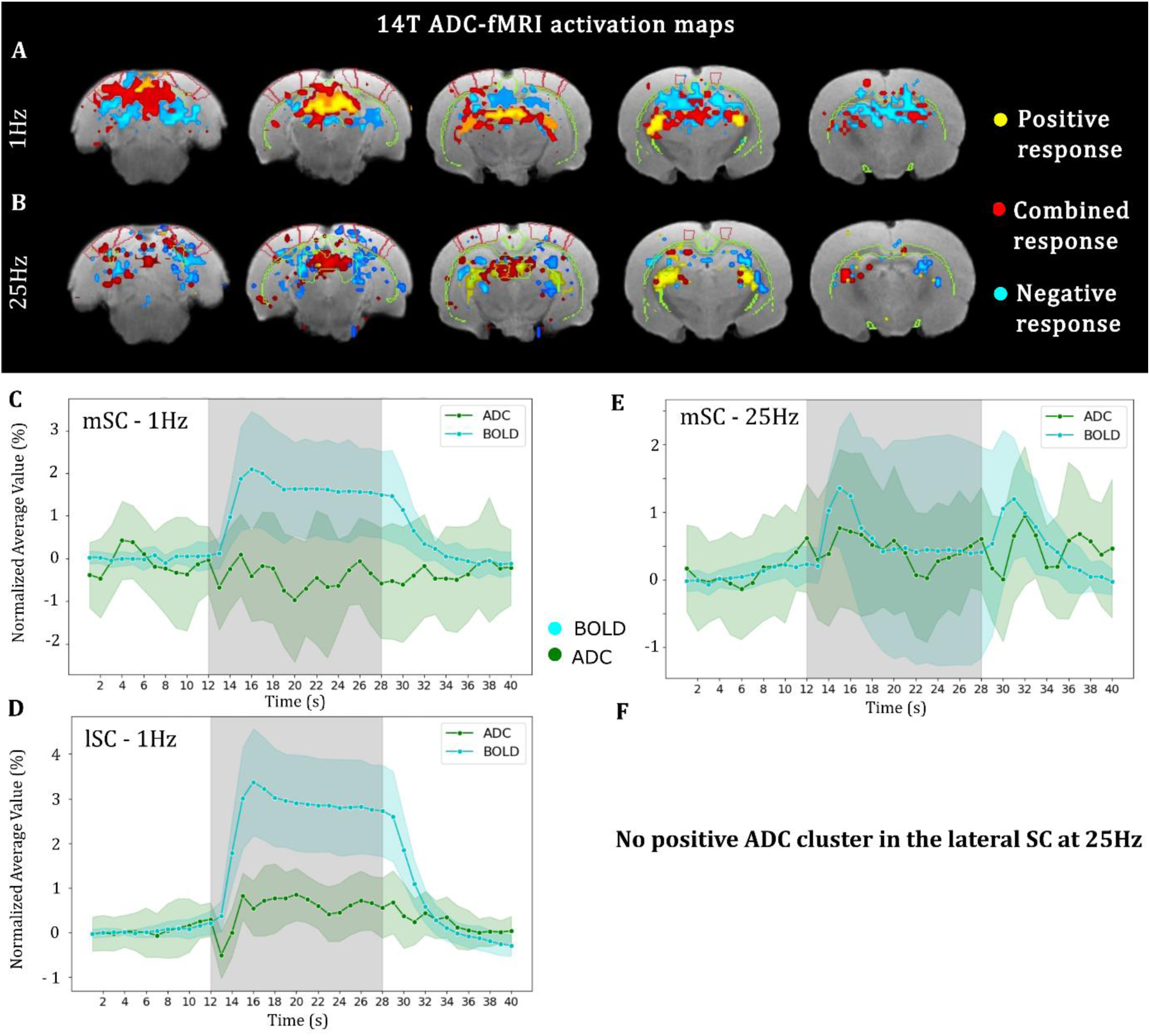
ADC response to 1Hz and 25Hz visual stimulation in comparison to BOLD at 14T. A-B: K-means clustered voxels with average response to (A) 1Hz and (B) 25Hz visual stimulation. C-F: Average response in the SC to (C-D) 1Hz and (E-F) 25Hz visual stimulation in (C & E) voxels with negative ADC response detected in the medial SC, and (D & F) voxels with positive ADC response detected in the lateral SC. Shading denotes standard deviation. Grey area represents stimulus duration. Higher vascular contribution than at 9.4T leads to activation maps more similar to BOLD. Only few voxels show a negative ADC response in the medial SC (C) at 1Hz with higher variability than at 9.4T.

